# MEG correlates of empty subjects in Japanese control and raising constructions

**DOI:** 10.64898/2026.01.08.698326

**Authors:** Koki Yamaguchi, Emi Yamada, Hiroshi Shigeto, Shinri Ohta

## Abstract

Empty categories are unpronounced elements with syntactic properties that play a central role in theories of sentence structure. Although there are several types within these categories, the neural basis for distinguishing among them remains unclear. Using magnetoencephalography, we investigate whether the brain distinguishes between two Japanese sentence structures: Control and raising. Although these constructions appear similar on the surface, they are argued to involve different types of empty categories. In theoretical analyses, the control-type empty category, which is called PRO, is often treated as an anaphoric element, similar to reflexives such as himself and herself, whereas the raising-type empty category is a noun phrase trace.

Twenty-six native Japanese speakers participated in a reading task under three experimental conditions: Control, raising, and baseline. Source estimates were computed, and condition differences were tested using spatiotemporal cluster-based permutation *t*-tests. We observed late left-hemispheric differences at approximately 700–800 ms after the critical verb. The control condition elicited larger responses than the raising condition, with activity centered in the temporal cortex spanning the middle temporal gyrus and the superior temporal sulcus and gyrus and extending into the anterior insula and the supramarginal gyrus. In addition, the control condition elicited larger late responses than the baseline condition in a broader left fronto-temporal distribution, including the inferior frontal cortex, anterior temporal cortex, and insula. These results provide source-level evidence that brain activity in the left language network differs between the control and raising conditions in Japanese during online sentence comprehension. Furthermore, they suggest that we can distinguish empty category types in the brain.

## Introduction

One key property of natural language processing is that comprehenders often interpret information that is not overtly expressed. For example, in “This is the book which I have read yesterday,” the fronted noun phrase “the book” is interpreted as being in a silent object position. Many frameworks treat this silent position as an empty category (EC).

Many neuroimaging studies have examined sentences that require interpreting an EC, including *wh*-dependencies, noncanonical word orders, and unpronounced arguments. Across these paradigms, activity is most often reported in a left-hemispheric fronto-temporal network [1–4]. The left inferior frontal gyrus (LIFG) has been shown to be a prominent frontal site. Many studies have also reported accompanying temporal involvement, especially in the superior and middle temporal regions. However, it remains unclear how broadly these findings can be generalized across different EC types.

One such case is unpronounced subjects in embedded infinitival clauses. Two superficially similar constructions are often distinguished here: control and raising. For example, “John tried to understand the formula” (control) and “John seemed to understand the formula” (raising). In both sentences, the embedded verb phrase “to understand the formula” appears without an overt subject in the main clause. Nevertheless, the intended interpretation is that Barnett is the one who understands. Despite this surface similarity, the two constructions are analyzed as involving different types of ECs. The control sentences contained PRO, an unpronounced subject. PRO is often described as an anaphoric element [5, 6] and its interpretation is constrained by an antecedent, much like reflexives such as himself or herself.

Raising sentences are often analyzed as involving a noun phrase (NP) trace. An NP-trace is a silent copy-like position associated with a noun phrase that is interpreted in more than one syntactic position. A familiar case is passive sentences. In a sentence “John was praised,” John is pronounced as the surface subject, but it is also interpreted as the understood object of “praise” in the corresponding active sentence (“Someone praised John”). The unpronounced object position can be treated as containing an NP-trace of Barnett.

Behavioral work across languages suggests that control and raising can differ during online comprehension [7–11]. However, direct neural evidence that clearly distinguishes the two remains limited. One influential ERP study reported a late difference between raising and control in the 600–800 ms time range [12].

However, this evidence is based on sensor-level ERPs and does not specify where in the brain the two constructions diverge. The present study addresses this gap by using magnetoencephalography (MEG) with source estimation to test whether control and raising elicit separable neural signatures during online sentence comprehension, and to localize the cortical regions that support the interpretation of an unpronounced embedded subject.

We focus on Japanese to test whether contrasts established largely in English also emerge in a language with a different basic word order. For example, English places the verb before its object (John read the book), whereas Japanese typically places the object before the verb (Taro-ga hon-o yon-da, lit. “Taro the book read”). Japanese has also been argued to distinguish control and raising predicates [13–16], and a recent behavioral study reported processing differences between Japanese control and raising constructions [10].

We compared three conditions: control, raising, and a baseline condition without an embedded infinitival clause and without an unpronounced embedded subject. We expected both control and raising to engage the fronto-temporal language network more than baseline, reflecting the additional computational demands required to interpret an unpronounced embedded subject. We further expected control and raising to show different spatiotemporal patterns within this network, because they differ in how the unpronounced subject is established and linked during comprehension.

## Materials and methods

### Participants

Twenty-six native Japanese speakers (13 males; mean age = 21.46 years, SD = 1.52) participated in this study. All participants showed right-handedness (mean laterality quotient = 93.4, SD = 16.3) as determined by the Edinburgh Handedness Inventory [17]. All participants had normal or corrected-to-normal vision and no history of linguistic, neurological, or psychiatric disorders. Written informed consent was obtained from all participants after a full explanation of the procedures. The study was approved by the Ethics Committee of the Graduate School of Medical Sciences at Kyushu University and conducted in accordance with the Declaration of Helsinki.

### Stimuli

Each stimulus sentence was divided into six frames. Frames 1–5 were identical across conditions, and Frame 6 was the critical frame because it contained the sentence-final compound verb that disambiguated the construction. The target materials comprised 90 experimental sentences (30 per condition: control, raising, baseline) and 90 filler sentences (180 sentences total) (Fig. 1A). Control and raising sentences in Japanese were implemented using sentence-final compound verbs [13]. Baseline sentences ended in a lexical compound verb and did not involve an embedded clause or an unpronounced embedded subject.

**Fig. 1.**
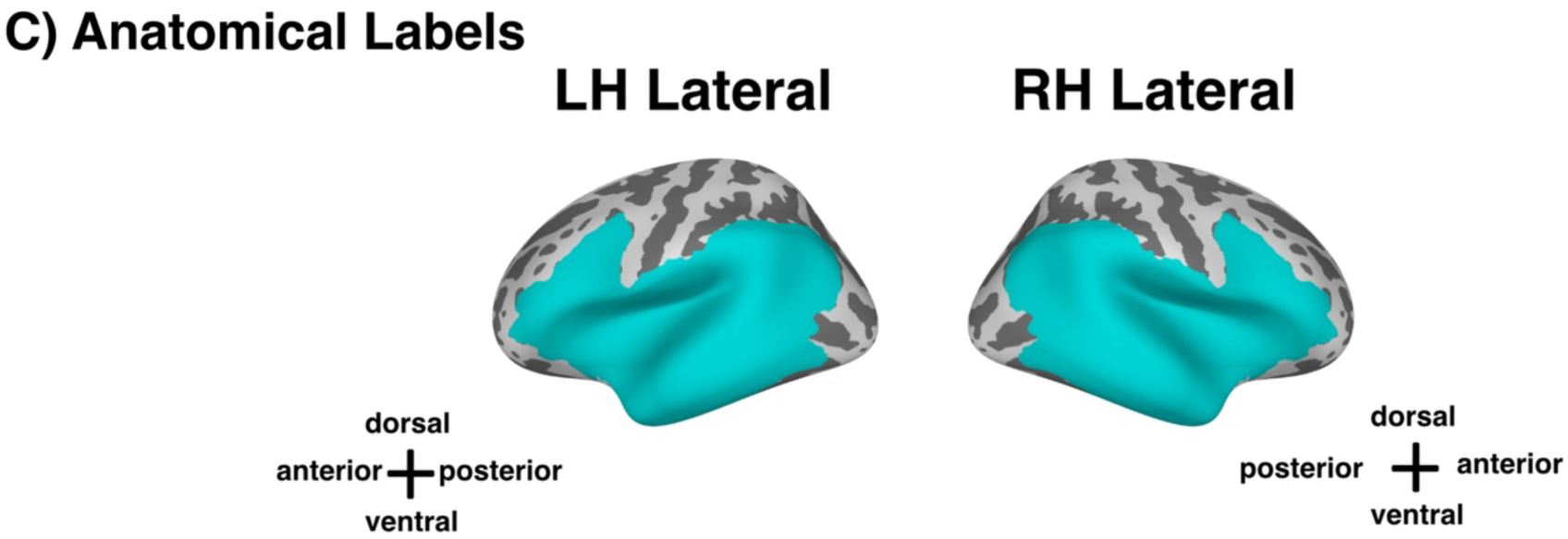
Overview of stimuli, procedure, and anatomical ROI. (A) Stimuli. Example Japanese sentences for the control, raising, and baseline conditions are shown from top to bottom. Each sentence is composed of six frames. For each condition, the first line provides the Japanese sentence in Hepburn romanization. Vowels with macrons indicate long vowels. The second line shows a word-by-word translation (gloss), and the third line provides an English translation. Empty categories are indicated at their theoretically assumed positions for illustration, but they were not presented to participants in experiments. (B) Procedure. Trial timeline showing the presentation duration for each region and the timing of the task. (C) Anatomical ROI. Labels defining the anatomical regions from the aparc. a2009s parcellation were used as the language-network ROI. Due to bioRxiv language policies, we excluded Panels A and B, which include example Japanese sentences. The original figure can be downloaded from the following link (https://doi.org/10.17605/OSF.IO/KNWRH).

To reduce the potential effects of the meanings of particular verbs, we used a wide range of verbs across control, raising, and baseline sentences; the full stimulus list is provided in the Supplementary Materials (https://doi.org/10.17605/OSF.IO/KNWRH). Lexical frequencies of the compound verbs were calculated using the Balanced Corpus of Contemporary Written Japanese (BCCWJ) [18] and the *Chunagon* web application (https://chunagon.ninjal.ac.jp/). We also confirmed that frequencies did not differ reliably across conditions using R (the “anovakun” function).

### Procedure

Prior to the main experiment, participants completed 10 practice trials that did not appear in the main session. These trials were used to familiarize participants with the task and to ensure that they understood the instructions.

During the MEG recording, participants were seated in an upright position approximately 2.5 m from a 32-inch LCD monitor (Display++; Cambridge Research Systems Ltd., Rochester, UK) inside a magnetically shielded room. The monitor had a refresh rate of 100 Hz. Stimuli were presented using PsychoPy (version 3.2.4; [19, 20]). A response mouse was placed on a table in front of the participant, and they were instructed to rest their fingers on the buttons throughout the experiment.

Each sentence was presented in six frames in Japanese script (*kanji* and *hiragana*).

Following previous research on ECs and antecedent reactivation[7, 10], participants performed a probe word recognition task after reading each sentence. Participants judged whether the probe word had appeared in the preceding sentence by pressing the left mouse button (“yes”) or the right mouse button (“no”). In target trials, the probe was the sentence-initial noun phrase. To discourage participants from predicting when and where the probe would appear, probe words were presented in filler sentences in various sentence frames.

Each trial began with a red fixation cross at the center of the screen. The six sentence frames were then presented sequentially with fixed exposure times, following our previous self-paced reading study using similar materials [10]. Each frame was followed by a brief white fixation and a short inter-stimulus interval. After the final frame, a red fixation cross was shown again, followed by the probe word, which remained on the screen until a response was made or for a fixed duration.

The experiment was divided into nine blocks, with short breaks every 20 trials to minimize fatigue. Before the sentence-reading task, a 3-minute eyes-open resting-state recording was acquired for the estimation of noise covariance.

### Data acquisition

Neuromagnetic activity was recorded using a whole-head 306-channel MEG system (Elekta Neuromag, Helsinki, Finland) comprising 102 magnetometers and 204 planar gradiometers. Signals were sampled at 1000 Hz with an online band-pass filter of 0.03–330 Hz. Four head position indicator (HPI) coils were attached to the scalp to monitor head position inside the dewar. A 3D digitizer (Fastrack; Polhemus, Colchester, VT, USA) was used to record the locations of three anatomical landmarks (nasion and the left and right preauricular points), the HPI coils, and additional points on the scalp surface. HPI measurements were obtained at the beginning of each recording session.

Vertical and horizontal electrooculograms were recorded using electrodes placed above and below the left eye and at the outer canthi of both eyes. Participants were instructed to minimize movements and to maintain fixation throughout the recording, and their posture was monitored with a video camera inside the magnetically shielded room. For source reconstruction, high-resolution T1-weighted structural MRI scans were acquired on a 3.0 T Achieva scanner (Philips N.V., Eindhoven, Netherlands).

### Pre-processing of MEG data

All MEG data were processed using MNE-Python version 1.4 [21]. Raw data were first denoised using oversampled temporal projection methods [22, 23]. Maxwell filtering [24, 25] was then applied to compensate for head movement and to remove external magnetic noise, and head positions were realigned to the position in the first recording session.

The sensor-level data were applied with a band-pass filter between 0.5 and 80 Hz and notch filtered at 60 Hz to remove line noise. Independent component analysis was used to identify and remove artifactual components related to eye blinks, eye movements, and cardiac activity. Continuous data were segmented into epochs from −200 to 1200 ms relative to the onset of the critical frame (Frame 6). Baseline correction was applied using the −200 to 0 ms pre-onset interval.

### Source estimation

Structural MRI data were processed with FreeSurfer [26] to reconstruct cortical surfaces and to generate three-shell boundary element method (BEM) head models using MNE-C tools [27]. Co-registration between MEG and MRI data was performed using the graphical interface of MNE-Python. Forward solutions were computed using the BEM surfaces, and source estimates were obtained with a unit-noise-gain linearly constrained minimum variance (LCMV) beamformer [28].

The noise covariance matrix was estimated from the 3-minute resting-state recording, and the data covariance matrix was computed from the 0–1200 ms post-onset window [29]. These covariance matrices were used to derive LCMV spatial filters, which were then applied to the event-related field data. For group analyses, individual cortical surfaces were morphed to the “fsaverage” template based on the MNI-305 standard brain provided by FreeSurfer. The source space was defined using an octahedral (oct-6) subdivision scheme, yielding 4098 vertices per hemisphere with a spacing of approximately 4.9 mm. All source estimates were computed at the whole-brain level and then restricted to predefined regions of interest for statistical testing.

### Statistical analysis

#### Behavioral analysis

Behavioral performance on the probe recognition task was analyzed using one-way repeated-measures analyses of variance (ANOVAs) with Condition (control, raising, baseline) as a within-participant factor. Analyses were conducted separately for response times (RTs) and accuracy. For RTs, only correct responses on target trials (90 sentences per participant) were included, and RTs were log-transformed to reduce skewness. ANOVAs were implemented in R (version 4.2.1) using the “anovakun” function (version 4.8.7; https://riseki.cloudfree.jp/?ANOVA%E5%90%9B).

#### MEG analysis

Statistical analysis of the source-level MEG data focused on differences in neural activity between conditions within a predefined time window. Nonparametric spatiotemporal cluster-based permutation tests were applied in order to identify clusters of vertices and time points showing reliable condition effects while controlling for multiple comparisons across space and time. In order to assess the magnitude and direction of observed effects, repeated-measures ANOVAs were performed with Condition (control, raising, baseline) as a within-participant factor. Analyses were implemented in R (version 4.2.1) using the “anovakun” function (version 4.8.7). When significant effects emerged, pairwise *t*-tests were conducted as post hoc comparisons, corrected for multiple testing using Shaffer’s correction.

Cluster-based permutation *t*-tests were performed separately for each pairwise comparison (control vs. raising, control vs. baseline, raising vs. baseline) within partially overlapped split time windows (300–700 and 600–1000 ms). We focused on the onset of the compound verb (Frame 6) (Fig. 1B). As the first five frames were identical across conditions, they were not included in the statistical analyses.

Analyses were restricted to an anatomically defined bilateral language network based on the aparc. a2009s parcellation [30]. This network included frontal, temporal, parietal, and insular regions in the left hemisphere (e.g., inferior frontal gyrus, precentral sulcus, superior and middle temporal gyri, supramarginal and angular gyri, and insula), as well as their right-hemisphere homologues (Fig. 1C).

## Results

### Behavioral data

Figure 2 summarizes the behavioral performance in the probe recognition task. Mean RTs for correct responses were comparable across the three conditions (Fig. 2A). Raw mean RTs were 870.0 ± 23.89 ms (mean ± SEM) for control, 881.3 ± 25.02 ms for raising, and 904.4 ± 23.88 ms for baseline. A one-way repeated-measures ANOVA on log-transformed RTs revealed no significant main effect of Condition (*F* (2,50) = 2.1927, *p* = 0.1222, *η^2^_p_* = 0.013). Thus, probe recognition speed did not differ reliably between control, raising, and baseline sentences.

**Fig. 2.**
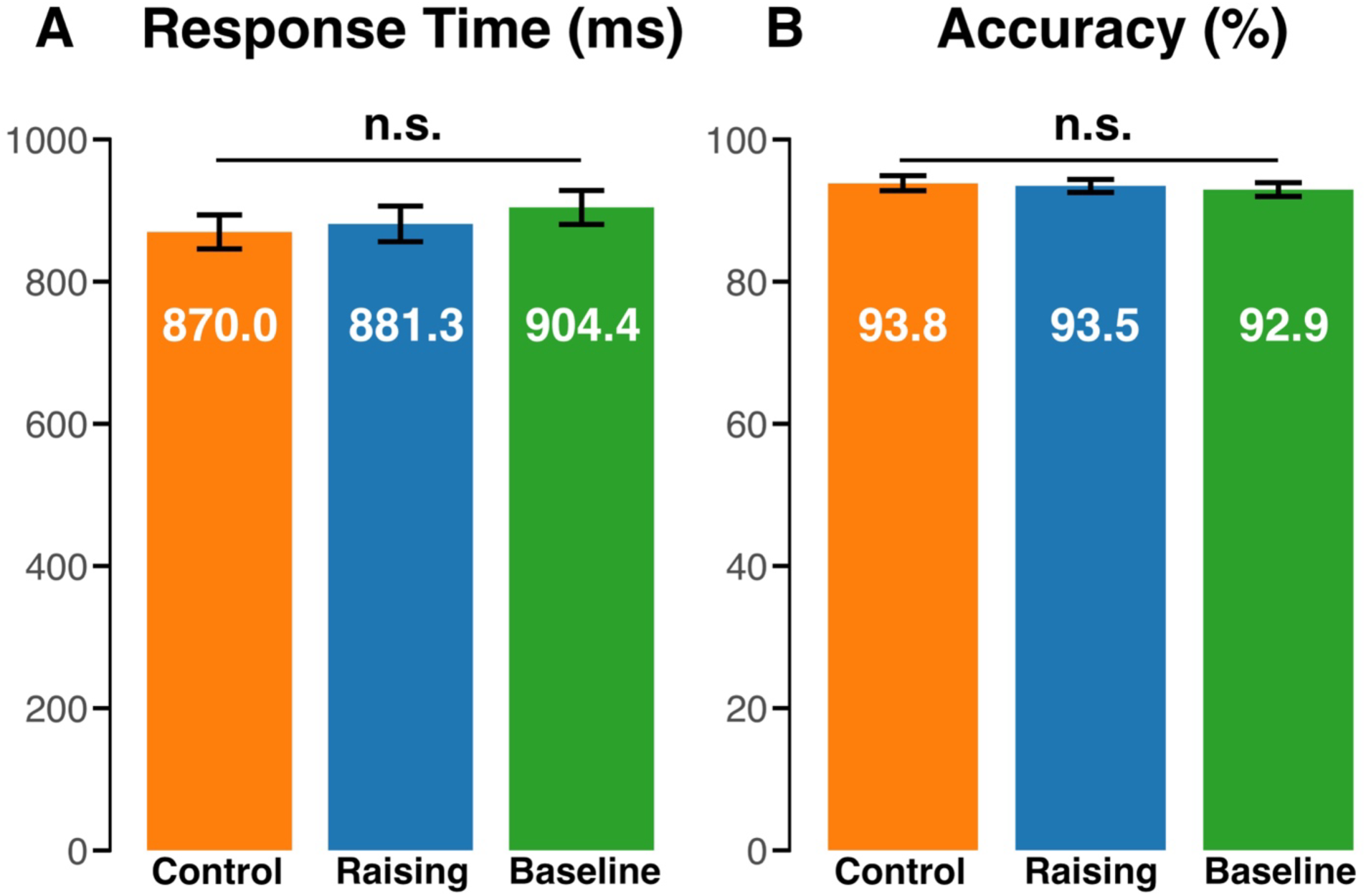
Behavioral results from the probe word recognition task. (A) Mean response times (RTs) to the probe word. (B) Mean accuracy. No significant effect of Condition was observed for either measure. Error bars indicate standard error of the mean (SEM). n.s.: non-significant.

Accuracy was high in all conditions, with mean values exceeding 90%. Raw mean accuracy was 93.8 ± 1.06 % (mean ± SEM) for control, 93.5 ± 0.91 % for raising, and 92.9% ± 0.97 % for baseline. The ANOVA on accuracy (Fig. 2B) likewise showed no significant effect of Condition (*F* (2,50) = 0.3802, *p* = 0.6857, *η^2^_p_* = 0.0056). These results indicate that participants performed the task accurately and that any neural differences between conditions cannot be attributed to differences in overall task difficulty or speed–accuracy trade-offs at the behavioral level.

### MEG data

#### Control vs Raising

A significant cluster (721–826 ms, *p* = 0.047, averaged *t* = 0.501) was identified in the left hemisphere (Fig. 3A) within the 600–1000 ms time window for the contrast between the control and raising conditions. In this cluster, activity was stronger for control than for raising. The cluster spanned the temporal cortex, extending across the middle temporal gyrus and superior temporal sulcus/gyrus (MTG, STS, STG), and further into the anterior insula and supramarginal gyrus. The largest differences were observed in MTG, STS, and STG.

**Fig. 3.**
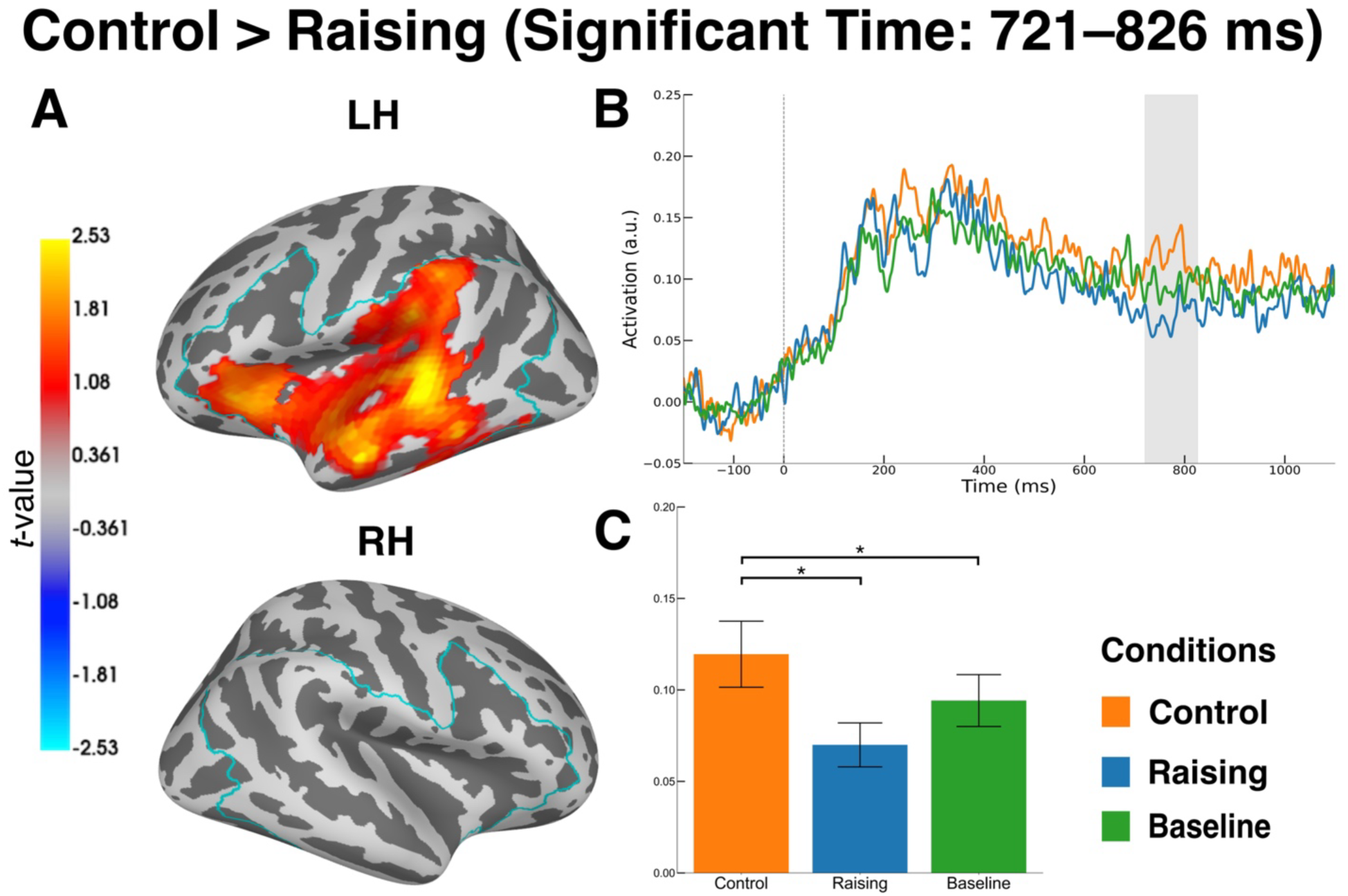
MEG results of permutation spatio-temporal cluster t-test: control vs. raising. (A) Significant cluster for the control > raising contrast, spanning the inferior parietal regions, the superior and middle temporal cortex, and the insula cortex. The color bar indicates t-values, averaged across the significant time window. Cyan outlines the ROI used for the statistical test. (B) Time course of activity averaged within the significant cluster, shown as condition-wise waveforms. The gray shaded region shows the significant time window. (C) Mean source activities across the significant cluster and significant time window. Asterisks indicate significant pairwise differences based on post hoc *t*-tests with Shaffer’s correction: *p* < .05 (*). Error bars indicate standard error of the mean (SEM).

The time course of activation within the cluster showed that the control condition was the largest compared to the other conditions (Fig. 3B and 3C). A significant main effect of Condition was observed (*F* (2, 50) = 6.4621, *p* = 0.0032, η²ₚ = 0.0682). Pairwise comparisons (Shaffer-corrected) showed that control was significantly greater than raising (*p* = 0.0161, *t* = 3.05) and baseline (*p* = 0.0294, *t* = 2.31), while no significant difference was found between raising and baseline (*p* = 0.0876, *t* = 1.78).

In contrast to this late left-hemispheric effect, no additional reliable differences between control and raising were observed in the early time window (*p* = 0.281, averaged *t* =0.322).

Similarly, in the right hemisphere, cluster-based tests revealed no significant clusters in the 300–700 ms window (*p* = 0.84, averaged *t* = 0.174) and the 600–1000 ms window (*p* = 0.179, averaged *t* = 0.476).

#### Control vs Base

For the comparison between control and base sentences, a significant cluster (710–811 ms, *p* = 0.045, averaged *t* = 0.806) was observed in the left hemisphere in the 600–1000 ms window (Fig. 4A). This cluster extended across the inferior frontal gyrus and sulcus, the insular cortex, the anterior MTG, STS, and STG. The time course of activation within the cluster showed that the control condition was the largest compared to the other conditions (Fig. 4B and 4C). A significant main effect of Condition was observed (*F* (2, 50) = 7.6188, *p* = 0.0013, η²ₚ = 0.1002). Pairwise comparisons (Shaffer-corrected) revealed that the control condition elicited significantly greater responses than baseline (*p* = 0.0015, *t* = 3.99) and raising (*p* = 0.0112, *t* = 2.74), while no significant difference was found between raising and baseline (*p* = 0.7281, *t* = 0.35). No significant clusters were detected in the early time window in the left ROI (*p* = 0.607, averaged *t* = 0.099) or in either time window in the right ROI (*p* = 0.953, averaged *t* = 0.104 for early time window; *p* = 0.273, averaged *t* = 0.28 for late time window).

**Fig. 4.**
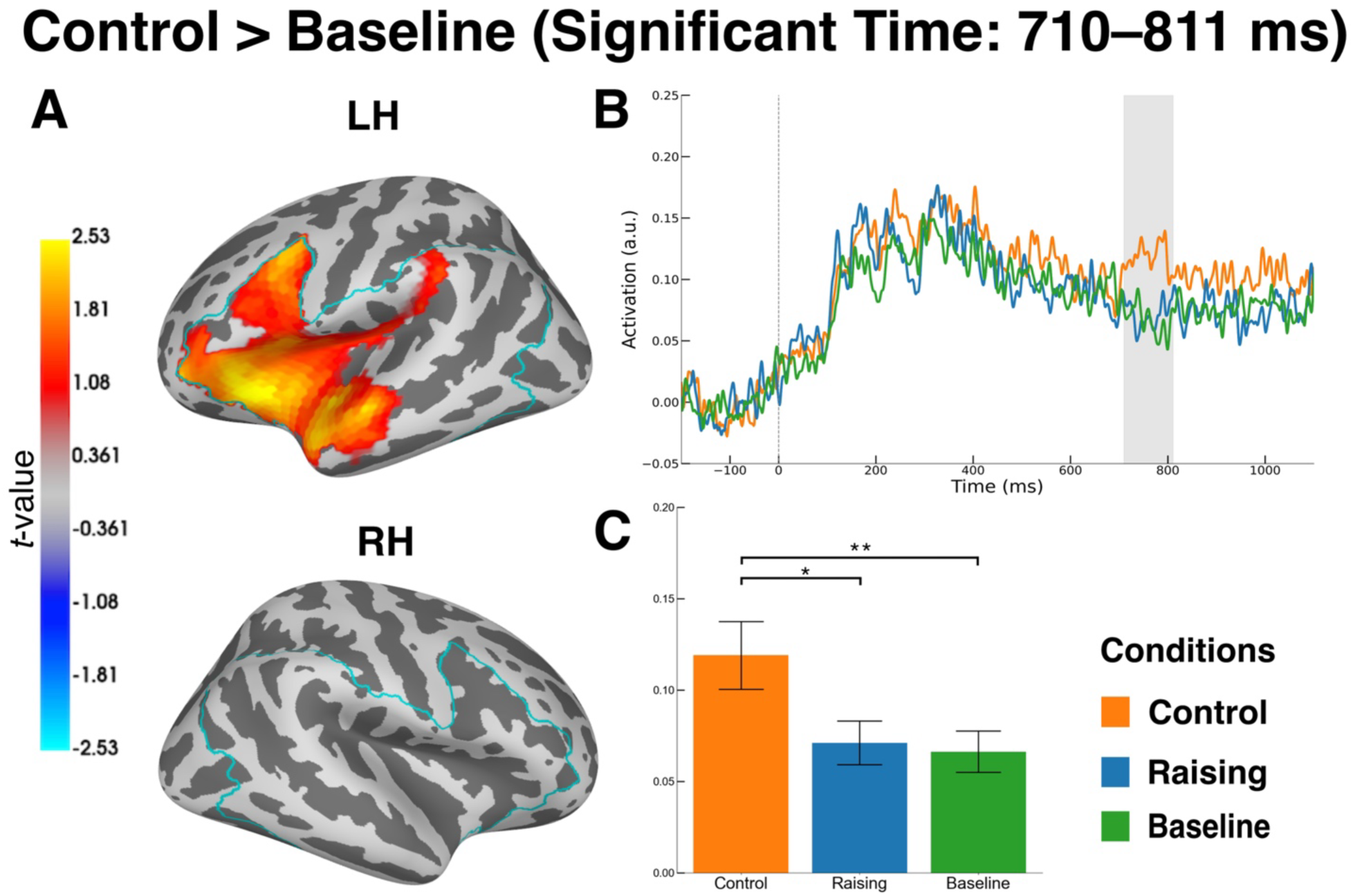
MEG results of permutation spatio-temporal cluster t-test: control vs. baseline. (A) Significant cluster for the control > baseline contrast, spanning the inferior frontal cortex, the insula cortex, and the anterior temporal regions. The color bar indicates t-values, averaged across the significant time window. Cyan outlines the ROI used for the statistical test. (B) Time course of activity averaged within the significant cluster, shown as condition-wise waveforms. The gray shaded region shows the significant time window. (C) Mean source activities across the significant cluster and significant time window. Asterisks indicate significant pairwise differences based on post hoc *t*-tests with Shaffer’s correction: *p* < .05 (*), *p* < .01 (**). Error bars indicate standard error of the mean (SEM).

#### Raising vs Base

For the comparison between raising and base sentences, a significant cluster (692–853 ms, *p* = 0.04, averaged *t* = −0.245) was observed in the left hemisphere in the 600–1000 ms window (Fig. 5A). This cluster was mainly across the middle temporal gyrus, superior temporal sulcus, and superior temporal gyrus. In a significant time window, baseline sentences elicited greater source-level activity than raising sentences (Fig. 5B and 5C). A significant main effect of Condition was observed (*F* (2, 50) = 6.5669, *p* = 0.0029, η²ₚ = 0.0699). Pairwise comparisons (Shaffer-corrected) showed that baseline elicited significantly greater responses than raising (*p* = 0.0083, *t* = 3.32), and that control was significantly greater than raising (*p* = 0.0307, *t* = 2.29).

**Fig. 5.**
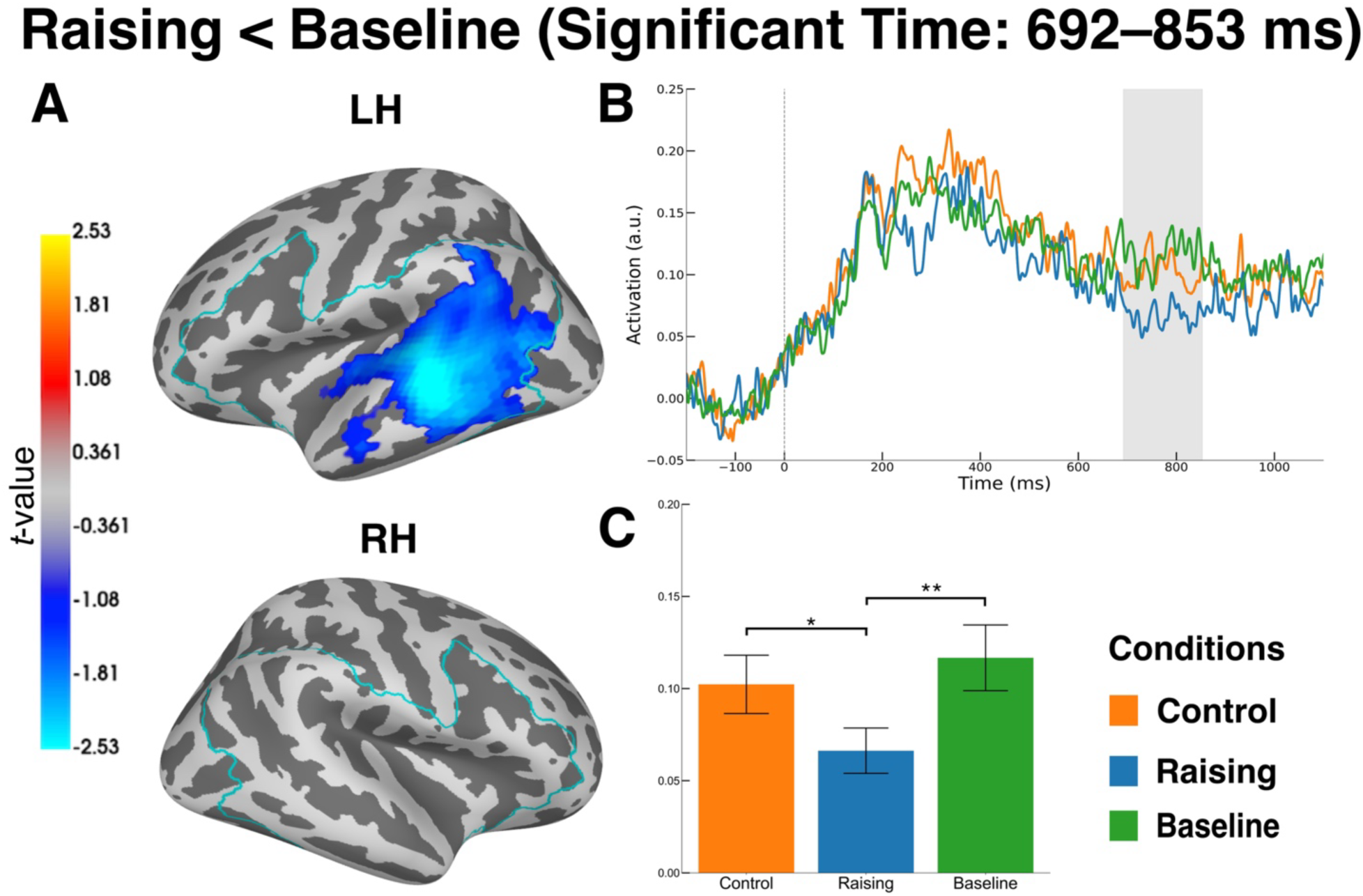
MEG results of permutation spatio-temporal cluster t-test: raising vs. baseline. (A) Significant cluster for the raising < baseline contrast, centered on the posterior superior and middle temporal cortex. The color bar indicates t-values, averaged across the significant time window. Cyan outlines the ROI used for the statistical test. (B) Time course of activity averaged within the significant cluster, shown as condition-wise waveforms. The gray shaded region shows the significant time window. (C) Mean source activities within the significant cluster and significant time window. Asterisks indicate significant pairwise differences based on post hoc *t*-tests with Shaffer’s correction: *p* < .05 (*), *p* < .01 (**). Error bars indicate standard error of the mean (SEM).

No significant difference was found between control and baseline (*p* = 0.2304, *t* = 1.23). No significant clusters were detected in the early time window in the left ROI (*p* = 0.633, averaged *t* = −0.218) or in either time window in the right ROI (*p* = 0.928, averaged *t* = −0.072; *p* = 0.844, averaged *t* = −0.194).

## Discussion

In this study, we investigated whether the contrast between control and raising —both involving EC— is reflected in neural activity during sentence comprehension. At the behavioral level, accuracy and response times in the probe task did not differ across conditions (Fig. 2), replicating our previous Japanese study using a similar paradigm [10]. In the MEG analysis, we observed late left-lateralized effects in both contrasts of interest. In the control–raising contrast, activity was stronger for control than for raising (Fig. 3A). The effect formed a cluster spanning the temporal cortex and insula. In the control–baseline contrast, activity was stronger for the control than for the baseline. The effect formed a left fronto-temporal cluster, spanning the inferior frontal, anterior temporal, and insular regions. Across the two control comparisons, we observed an overlap in the appearance of late clusters. Both the control vs. raising and control vs. baseline contrasts extended into an anterior temporal–insula region, indicating that this area was engaged in both contrasts when control was compared with the other conditions (Figs 3A and 4A). Prior work on anaphoric dependencies has linked temporal regions to the establishment of an anaphor–antecedent relation [31–34]. Based on this, the overlapping anterior temporal activity in our control contrasts suggests that control may place greater demands on processes related to establishing a referential dependency for an unpronounced subject. Therefore, we propose that the shared anterior temporal–insula effect may partly reflect computations relevant to the anaphoric properties of PRO in control sentences.

Furthermore, the timing of our effects provides additional constraints. The shared effects in the anterior temporal cortex and insula emerged in a late window around 700–800 ms (Figs 3B and 4B). This timing aligns with prior work comparing control and raising [12], which reported a 600–800ms time window difference. It is also in line with ERP studies on anaphoric dependencies that report late positivities in a similar window during anaphor resolution [35, 36].

Finally, our results did not support the predicted increase for raising relative to baseline (Fig. 5A). We expected raising to elicit stronger responses than baseline because raising includes an embedded infinitival clause and an empty subject position, often analyzed as an NP-trace.

This pattern did not emerge. Instead, baseline showed stronger temporal activation than raising, as shown in Fig. 5C. We also expected greater engagement of the left-lateralized language network centered on LIFG for raising, but we did not observe this pattern. Two factors may have contributed to these results. The first factor was predictability. In much prior work on ECs, the antecedent precedes the empty position, so comprehenders can anticipate that a dependency will need to be completed later. For example, in a sentence “Which book did you buy yesterday?,” when a *wh*-phrase such as “which book” appears at the beginning of a sentence, the EC can be expected later at the verb “buy.” Some studies have suggested that LIFG responses, often linked to EC processing, may partly reflect cue-based prediction [37, 38]. In our materials, by contrast, the relevant empty-subject analysis becomes clear only at the sentence-final compound verb.

Thus, comprehenders must process the compound verb before the type of empty subject and its antecedent can be identified. The existence of EC cannot be predicted in advance. This feature of our design may have reduced the chance of observing a raising-over-baseline increase. The second factor concerns the baseline condition. Although baseline sentences are structurally the simplest because they lack an embedded clause and any empty subject, they still require substantial lexical–semantic computation to compose the meaning of the compound verb. Under this view, reduced demands from dependency-related operations may allow greater engagement of temporal regions that support lexical access and integration during sentence comprehension.

Of course, this logic also allows the possibility that baseline could elicit stronger responses than control in temporal regions. In the present data, however, the reliable cluster in the control–baseline comparison was in the opposite direction (control > baseline). This pattern suggests that, in our materials and time window, processes required to establish and integrate the PRO dependency in control imposed additional demands that exceeded the lexical–semantic composition demands of the baseline compound verb, resulting in a larger late response for control.

A limitation of the present study is that MEG source estimates are correlational and therefore cannot establish which computations causally generate the observed effects.

Accordingly, although the larger responses for control are consistent with prior work linking temporal regions to the formation of an anaphor–antecedent relation [29–32], the interpretation that our control effects reflect processes related to this relationship remains indirect and requires further support. Future work can strengthen this interpretation by providing a more detailed characterization of the underlying neural dynamics (e.g., through time–frequency analyses) and by complementing the correlational evidence from MEG with causal approaches, such as transcranial electrical stimulation, to test whether the implicated regions and dynamics are necessary for sentence comprehension.

## Conclusion

In this study, we tested whether Japanese control and raising constructions can be distinguished in MEG during sentence comprehension. Control sentences elicited larger late responses than both raising and baseline sentences in a left-lateralized network centered on the temporal cortex and insula. These results show that superficially similar sentences with an unpronounced embedded subject can evoke separable neural signatures, providing source-level evidence that the left language network is sensitive to construction-specific mechanisms for interpreting ECs.

## CRediT authorship contribution statement

Koki Yamaguchi wrote the first draft of the manuscript. All authors contributed to the conception and design of the study, prepared the task materials, performed the statistical analysis, and revised the manuscript.

## Acknowledgments

We would like to thank all the members of our laboratory. We would also like to thank Taira Noriko for her administrative support.

## Data availability

The data are available upon request to the corresponding author.

